# Neural network for the prediction of treatment response in Triple Negative Breast Cancer ^*^

**DOI:** 10.1101/2022.01.31.478433

**Authors:** Peter Naylor, Tristan Lazard, Guillaume Bataillon, Marick Lae, Anne Vincent-Salomon, Anne-Sophie Hamy, Fabien Reyal, Thomas Walter

## Abstract

The automatic analysis of stained histological sections is becoming increasingly popular. Deep Learning is today the method of choice for the computational analysis of such data, and has shown spectacular results for large datasets for a large variety of cancer types and prediction tasks. On the other hand, many scientific questions relate to small, highly specific cohorts. Such cohorts pose serious challenges for Deep Learning, typically trained on large datasets.

In this article, we propose a modification of the standard nested cross-validation procedure for hyper-parameter tuning and model selection, dedicated to the analysis of small cohorts. We also propose a new architecture for the particularly challenging question of treatment prediction, and apply this workflow to the prediction of response to neoadjuvant chemotherapy for Triple Negative Breast Cancer.

## 1 Introduction

### 1.1 Context

Breast cancer (BC) is the most common cancer in women and the leading cause of cancer deaths with 18.2% of deaths among female cancer patients and 8% among all cancer patients [1]. Out of the four main breast cancer types, Triple Negative Breast Cancer (TNBC) represents 10% of all BC patients. This group has the worst prognostic with a five-year survival rate of around 77 percent versus 93 percent for the others. Currently no specialised treatments exists and the standard procedure consists in administrating neoadjuvant or adjuvant chemotherapy [2]. TNBC research is still a very active field of study [3] and on the one hand, most works have focused on stratifying cohorts based on molecular and biological profiles [4]. We, on the other hand, tackle the problem of predicting the response variable in a TNBC neoadjuvant chemotherapy (NACT) cohort from a histological needle-core biopsy section from the primary tumour prior to treatment. We know that this is a relevant task as pathologist routinely check biopsy’s in order to derive treatment and diagnostic. In contrast to most of the effort in cancer research, which is driven by the analysis of sequencing data, our study is based solely on the histological image data prior to treatment.

Each histological sample corresponds to tissue slides encompassing the tumour and its surrounding, stained with agents in order to highlight specific structures, such as cells, cell nuclei or collagen. The morphological properties of these elements and their spatial organisation have been linked to cancer subtypes, grades and prognosis. Even if pathologists have been trained to understand and report the evidence found in this type of data, the complexity, size and heterogeneity found in histological specimens make it highly unlikely that all relevant and exploitable patterns are known of today. Tissue images are informative about morphological and spatial patterns and are therefore inherently complementary to omics data.

Two major technological advances have triggered the emergence of the field of Computational Pathology: first, the arrival of new and powerful scanners replaced to some extent the use of conventional microscopes. Today, slides are scanned, stored and can be accessed rapidly at any moment [5]. This in turn has lead to the generation of large datasets that can also be analyzed computationally. The second element was the rise of new computer vision techniques.

Indeed, while the analysis of tissue slides has been of interest to the Computer Vision community for many years [6], it is the advent of deep learning that has truly impacted the field. The advent of deep learning has stemmed a wide number of projects and investments. For visual systems, it is the combination of very large annotated dataset [7], hardware improvement and convolutional neural networks (CNN) that led to human-like capabilities. These outbreaks in performance have led to the creation of many annotated datasets and to the application of CNN’s to many tasks. However, for biomedical imaging in particular the application of CNN is not always straightforward:

1. The price for generating and annotating large biomedical dataset limits the progress of big data in this domain [8].
2. In histopathology in particular, each individual sample can be very large, one sample can be up to 60GB uncompressed. This leads to multiple issues, again linked with the time and price needed for annotation, but also for the subsequent analysis where ad hoc methods have to be used as an entire image does not comfortably fit in RAM.
3. The nature of the data is inherently complex, each biological sample has its own individual patterns to be differentiated with relevant pathological evidence. For histopathology data, samples have a very large inter-slide, but also intra-slide variability that make the apparent signal harder to detect. In addition, the level of detail can be an additional difficulty: the relevant image features may be very fine grained, such as mitotic events, or very large such as the size of relevant image regions (necrotic, tumorous) [9].

In this paper, we apply deep learning models to histopathology data, in particular a TNBC dataset with less than 350 slides. This context poses many difficulties, especially in terms of validation where we have to make the most out of our available data. We propose a more suitable validation procedure, prove its validity and efficiency with a simulation case study and finally apply it to our TNBC cohort. For this, we also present a new architecture and provide a benchmark with respect to other currently used methods.

The paper is organised as follows: in the next Section 1.2 we describe related work. In Section 2, we describe the methodological developments. In Section 2.1, we present the limits of the current validation procedures and our alternative method used in this study. We then introduce our histopathology dataset in Section 2.3. Section 2.4 is devoted to introducing the DNN architectures which will be applied to the TNBC cohort. In Section 3, we show our results on the simulated data for our validation procedure and the application of our DNN to the TNBC cohort. Finally in Section 4 we discuss our methods and results.

### 1.2 Related work

#### 1.2.1 Challenges in Computational Pathology

The main fields of research in computational pathology can be divided in three categories:

1. Preprocessing, in particular color normalisation which aims at reducing the bias introduced by staining protocols used in different centers [10, 11].
2. Detection, segmentation and classification of objects of interest, such as region [12, 13] and nuclei [14, 15].
3. The prediction of slide variables, such as presence of disease [16,17], survival [18,19], gene expression [20,21], genetic mutations [22] or genetic signatures [23, 24].

Pipelines for slide variable predictions are usually divided into several steps. Tiles are partitioned into smaller images, usually referred to as patches or tiles, which are then encoded by a DNN, often trained on ImageNet [18, 19], as depicted in Figure 2. The training on ImageNet might be surprising at first sight, as the nature of the images are very different. In addition, ImageNet samples usually have a natural orientation, where the main object of interest is usually centered and scaled to fit in the image [25]. In contrast, histopathology images have a rotationally invariant content with no prior regarding scale or positioning of the relevant structures. However, rotational invariance can be imposed [26, 27], and in practice ImageNet based encodings are widely used and tend to perform very well.

After encoding of all tiles, each WSI is converted into a *P* × *n*_*i*_ matrix where *P* is the encoding size and *n*_*i*_ the number of tiles. The last step consists in aggregating tile level encodings to perform unsupervised or supervised predictions at the slide level [17–19, 26, 28].

Computational Pathology as a field has benefited from the generation of large annotated data sets, mostly with pixel-level annotations [14, 16, 29], or cell-level annotations [30] for cell classification. The major resource for WSI with slide level annotations are the Cancer Genome Atlas (TCGA) and the Camelyon Challenge [16]. These public repositories are paralleled by many in-house datasets (thus not accessible to the public), some of which can be very large, namely in a screening context, e.g. [17]. In most cases however, the datasets tend to be very small and fall therefore in the *small n large p category*. This is due to the fact that often the most interesting studies focus on particular molecularly defined cancer subtypes for which only small cohorts exist. In addition, collecting the output variable might be very challenging and time-consuming, if the project is not formulated in the context of Computer Aided Diagnosis. This is particularly true for treatment response prediction.

#### 1.2.2 Challenges in applying DNN to small *n* large *p*

For all supervised learning method it is custom to use a two step procedure for estimating the performance. After dividing your dataset into three categories: train, validation and test. The first step consists in performing model selection with the training and validation set. The second step simply involves evaluating the chosen model on the test set in order to assess an unbiased estimator of the performance [31]. This is however only possible if the three categories are large enough. When the validation set is too small and the discrepancy in the data too high, one could very easily over-fit or under-fit on the dataset [32]. When the number of samples *n* is small, which is usually the case for biomedical data, alternatives validation methods have to be found such as cross-validation (CV) and nested cross validation (NCV). CV is mostly used for model selection or assessing performance. NCV is used when the model needs a tuning based on an external dataset, such as hyper parameter tunning for Support Vector Machines. Even if these methods have been debated [32–34], they are widely accepted. These methods are explained in more details in Section 2.1.

#### 1.2.3 Prediction of the response to neoadjuvant chemotherapy in TNBC

Neoadjuvant chemotherapy (NACT) responses varies among patients in TNBC and no clear biological signal has been shown. Survival in these cohorts have been correlated to the Residual Cancer Burden (RCB) variable [35] which can be used as a proxy for response. RCB is a pathological variable based on measurements of how much the primary tumour has shrunk and of the size of metastasis in axillary lymph nodes. Finding biological evidence to NACT response would allow for adequate and specific treatment, some histological variables have been found to be correlated with survival, such as the number Ki-67 positive cells [36], tumor infiltrating lymphocytes [37] and the Elston and Ellis grade [38]. Depending on the context, some alternative treatments have been found to help overall survival, such as those based on anthracycline and taxanes [2], carboplatin [39] or with olaparib and talazoparib [40]. Some treatments have emerged with targeted immunotherapy in combination with atezolizumab (anti-PD-L1 antibody) and nanoparticle albumin-bound (nab)-paclitaxel [40]. Most of the studies for NACT responses have been performed in clinical practices and based on pathological variables [36, 41, 42]. In addition, some studies have analysed sequencing and molecular profiles in order to better understand and stratify cohorts [4, 43, 44].

To the best of our knowledge, it remains unclear whether and to which extent NACT response can be predicted from biopsies taken prior to treatment, and only few works have addressed this question so far [26, 45].

## 2 Materials and Methods

### 2.1 Validation procedure

Here, we present our procedure that replaced NCV in order to train DNN in a context of *small n large p*. We first explain cross-validation (CV), nested cross-validation (NCV) and their limitations. We, propose a different procedure, better suited and show its effectiveness on a small case study.

#### 2.1.1 Cross-validation

CV is a common procedure for model evaluation or model selection, specifically in situations where the data set is relatively small. CV divides the initial data set into *k*_*f*_ folds, denoted 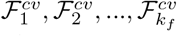 and runs algorithms on the data sets with one fold left out. We define, for all *j*, the set 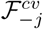, which is the union of all folds expect for fold *j*:

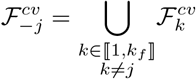

##### Cross validation for model selection

CV can be used for model selection or model tuning. The procedure that returns a tuned model ℳ will be notated *f*^*cv*^.

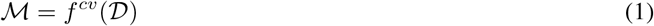

We give the pseudo code in Algorithm 1, where ℋ = *h*_1_, …*h*_*i*_, .. is the set of hyperparameters (HP).

###### Algorithm 1: Model selection, *f*^*cv*^

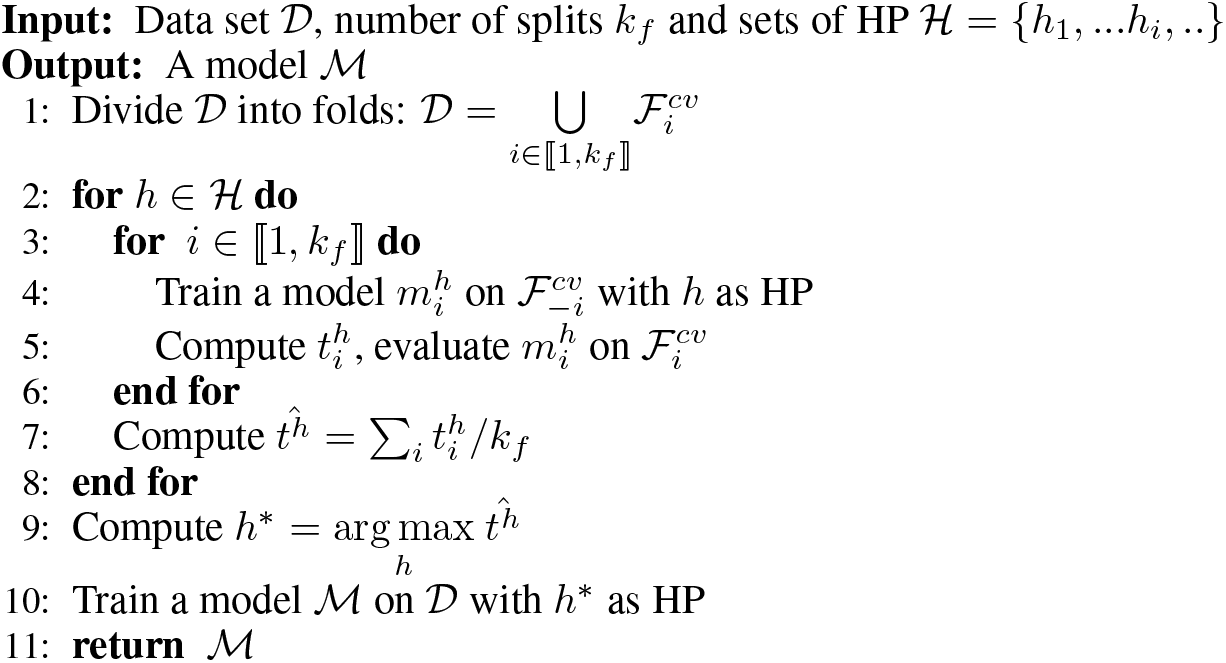

##### Cross-validation for model evaluation

We can use CV to evaluate a given model and a HP set *h*. The procedure is similar to the pseudo code given in Algorithm 1, however we give in input a model and only one hyperparameter set and return 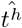. In this case, 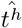 is an unbiased estimator of the performance, as no optimization of the hyperparameters took place. If however several sets of hyperparameters are tested as to minimize the accuracy measured by cross validation, this accuracy is an over-optimistic estimation of the true accuracy. In order to get a realistic estimation of the accuracy, we therefore have to turn to nested cross validation.

##### Nested cross-validation

NCV is a procedure that allows one to tune a model and effectively report an unbiased estimation of the performance of the tuned model.

Given sets of HP and a data set 𝒟, NCV corresponds to two nested loops of CV: The outer CV loop is for model evaluation, usually applied on test folds, sometimes referred to as outer folds. The inner CV loop is for model tuning. I.e. for each test fold, we perform a complete CV on the remaining data to correctly tune the model, and test the performance of the tuned model on data that has neither been used for training nor for HP tuning. We show the pseudo-code for NCV in Algorithm 2.

###### Algorithm 2: Nested cross validation

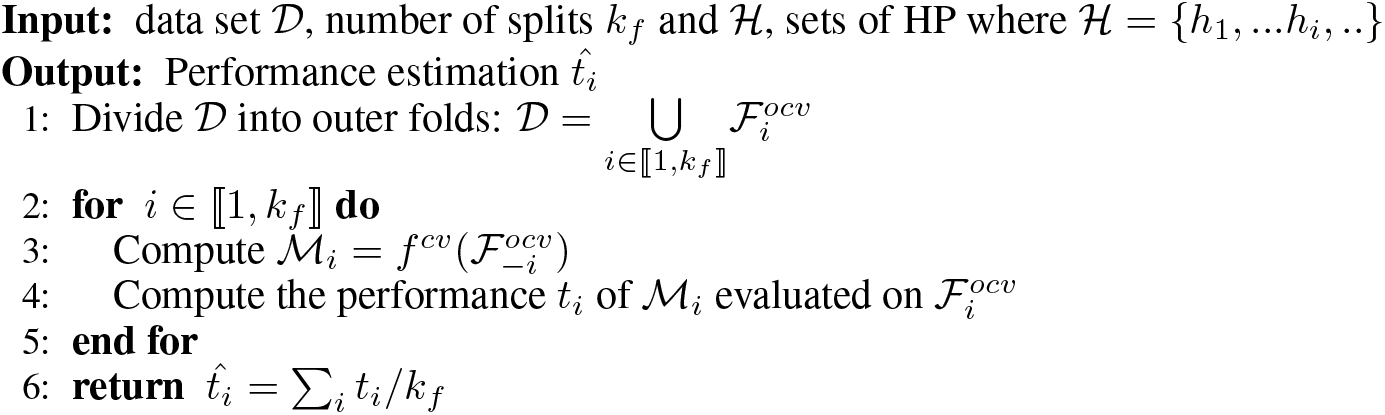

Another possible view is to see NCV as a simple CV for a model selection algorithm. For NCV, the model selection algorithm would be *f*^*cv*^.

It is important to note that as we are training DNN, we do not use a fixed hyperparameter set, ℋ, but randomly generate the set as it has been shown that randomised search performs better [46].

#### 2.1.2 Limitations

DNN training suffers from inherent randomness, as the loss function is highly non-convex and possess many symmetries [47]. In addition, there are some stochastic differences between different training runs, such as the random initialization of the weight parameters and the data shuffling, naturally leading to different solutions. Especially for small datasets, these stochastic variations lead to notable differences in performance when we repeat training with the same hyper-parameters.

In the classical setting, CV provides us with a set of hyper-parameters that lead to a model with optimal performance, as estimated in the inner loop. For DNN trained on small datasets, there is no guarantee that the same set of hyper-parameters will lead to similar performance, and for this reason retraining is not guaranteed to lead to a very good solution.

Another problem with the retraining in line 10 in Algorithm 1 is the use of early stopping. Early stopping is a very powerful regularization procedures that choses experimentally the point between the under- and over-fitting regime, but for this it requires a validation set. Early stopping would therefore not be applicable in the traditional CV-scheme with retraining.

#### 2.1.3 Nested ensemble cross validation

Due to the incompatibility between NCV and early stopping we propose to modify the model selection procedure, i.e. function *f*^*cv*^ shown in Algorithm 1. In particular we do not perform retraining and return an ensemble of the models used during CV. Similarly to NCV, we perform a CV where we propose to modify *f*^*cv*^ into a better suited procedure, named *f*^*ecv*^ (ensemble cross validation), shown in Algorithm 3.

##### Algorithm 3: Model selection, *f*^*ecv*^

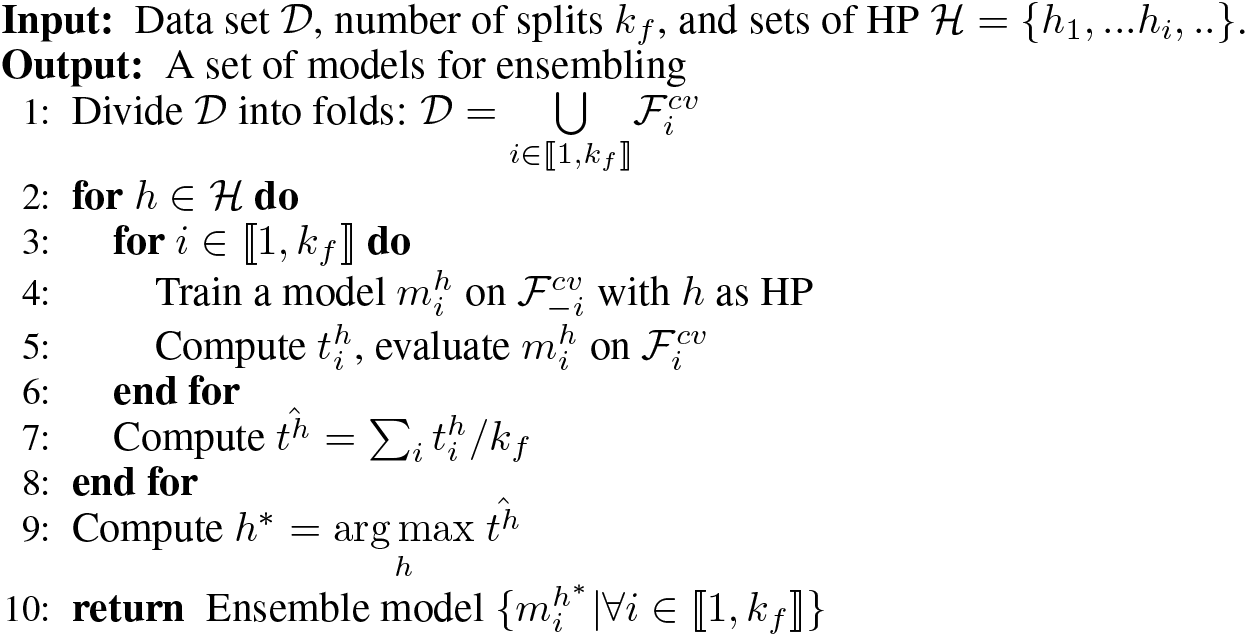

The main difference between *f*^*cv*^ and *f*^*ecv*^ is that we remove the final model retraining, i.e. line 10 of Algorithm 1 and give back the full set of *k*_*f*_ models trained for all folds for the maximizing hyperparameters; the prediction is obtained by ensembling of these models.

The advantage of this procedure is that we omit the retraining step which allows us to use early stopping for all individual models. In addition, we add another level of regularization by the ensembling. Of note, *f*^*ecv*^ can be used in an inner loop, too. This then leads to Nested Ensemble Cross Validation (NECV).

### 2.2 Simulations

In order to compare and validate our procedure presented in Section 2.1 we propose to conduct a series of simulation studies where the results will be given in Section 3.1. In particular, we wish to demonstrate that DNN training given a set of HP can lead to inconsistent models and that NCV therefore might provide under-performing models compared to NECV, the validation procedure we propose.

#### 2.2.1 Data set simulation

We simulated a simple balanced binary data, of size *N* = 350 in a medium to high dimensional setting with *p* = 256. We have ∀*i* ∈ ⟦1, *N* ⟧, *Y*_*i*_ ∈ {−1, 1} and 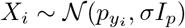 where *p*_*j*_ is a cluster center, *σ* a given standard deviation and *I*_*p*_ the identity matrix of size *p*. We set one cluster center at *p*_1_ = (1, 1, …, 1) and the second cluster center at *p*_−1_ = (−1, −1, …, −1). In Figures 1.A, 1.B, 1.C and 1.D respectively, we show four plots of the simulated data reduced to two dimensions thanks to a UMAP [48], with standard deviations set to 2, 6, 10 and 14 respectively. Naturally, when the standard deviation increases the data becomes less separable.

**Figure 1:**
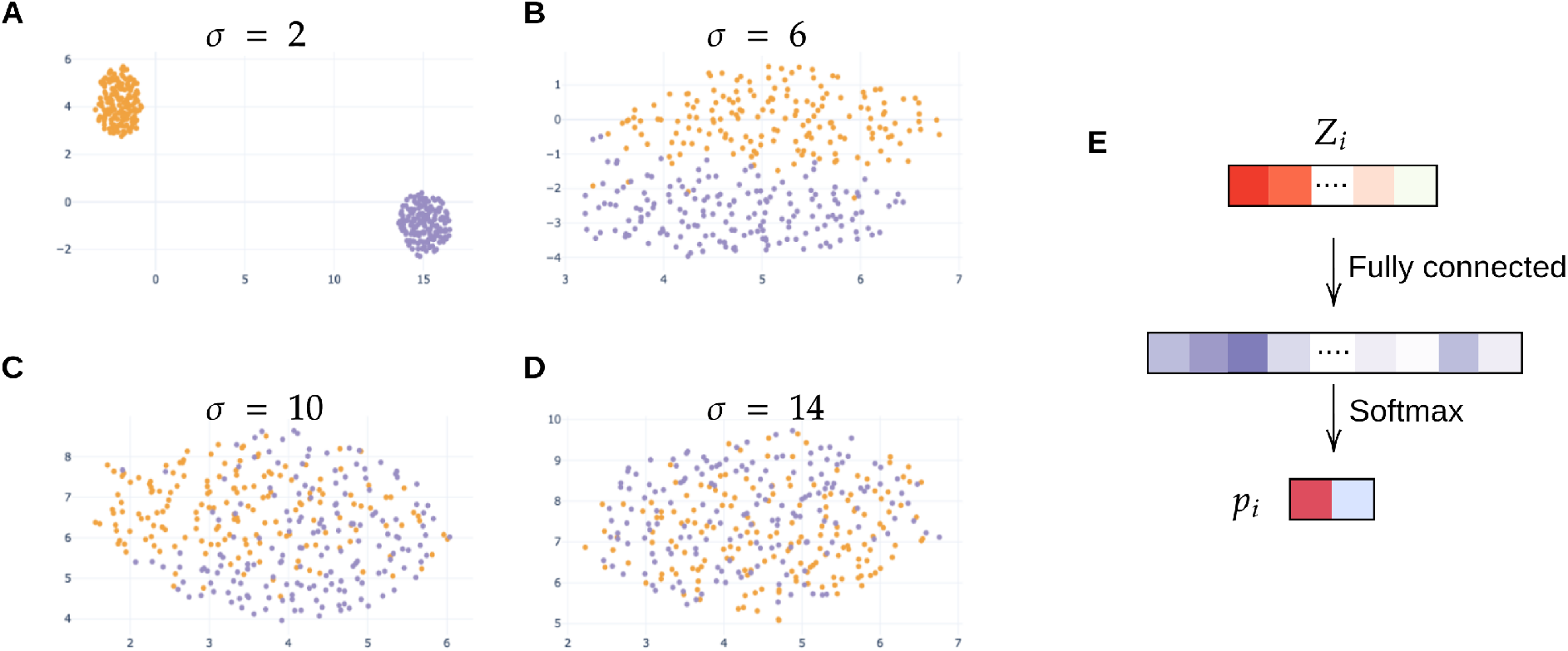
Data set simulation with *p* = 256 and model, (**A**) *σ* = 2, (**B**) *σ* = 6, (**C**) *σ* = 10, (**D**) *σ* = 14. (**E**) Simple two layer DNN model for predicting the class label.

#### 2.2.2 Model

We apply a simple DNN model, composed of two layers, a first hidden layer with 256 hidden nodes and a classification layer with 2 nodes, the model is depicted in Figure 1.E.

In particular, we minimise a cross-entropy error with an Adam optimiser and a batch size of 16. The tunable parameters are the weight-decay, drop out and learning rate. As an extra regularisation we use batch normalisation.

### 2.3 Application to histopathology data

#### 2.3.1 Data generation and annotation

The data set used was generated at the Curie Institute and consists of annotated H&E stained histology needle core biopsy sections at 40× magnification sampled from a patient suffering from TNBC. In this paper, we evaluate the prediction of the response to treatment based solely on a biopsy sectioned prior to the treatment. As discussed in the introduction, not all patients respond to NACT, and we are therefore aiming at predicting the response to NACT based on the biopsy. In particular, each section was quality checked by expert histopathologists.

For each patient, we also collect WSI after surgery, allowing an expert pathologist to establish the residual cancer burden, as a proxy for treatment success. Out of the 336 samples that populate our data set, 167 were annotated as RCB-0, 27 as RCB-I, 113 as RCB-II and 29 as RCB-III. This data set is twice as large as the data set used in our previous study [26]. Similarly to this study, we refine the number of classes in order to avoid the problem of under-represented class. We investigate two prediction settings: (1) pCR (no residuum) vs RCB (some residuum) and (2) pCR-RCB-I vs RCB-II-III, which is clinically more relevant, as it is informative of a patient’s prognosis.

#### 2.3.2 Data encoding

As each biopsy section is relatively big, we wish to reduce the computational burden of feeding the entire biopsy to our algorithms. Instead, given a magnification factor, we divide each biopsy into tiles of equal sizes, 224 × 224 and project this tile into a lower dimensional space. We use a pre-trained DNN on ImageNet [7] such as ResNet [49] which produces a encoding of size 2048. This process is illustrated in Figure 2 where each biopsy section is converted into a encoded matrix of size *n*_*i*_ × *P* where *P* is the size of the resulting encoding and *n*_*i*_ the number of tiles tissue extracted from tissue *i, i* ∈ ℕ.

**Figure 2:**
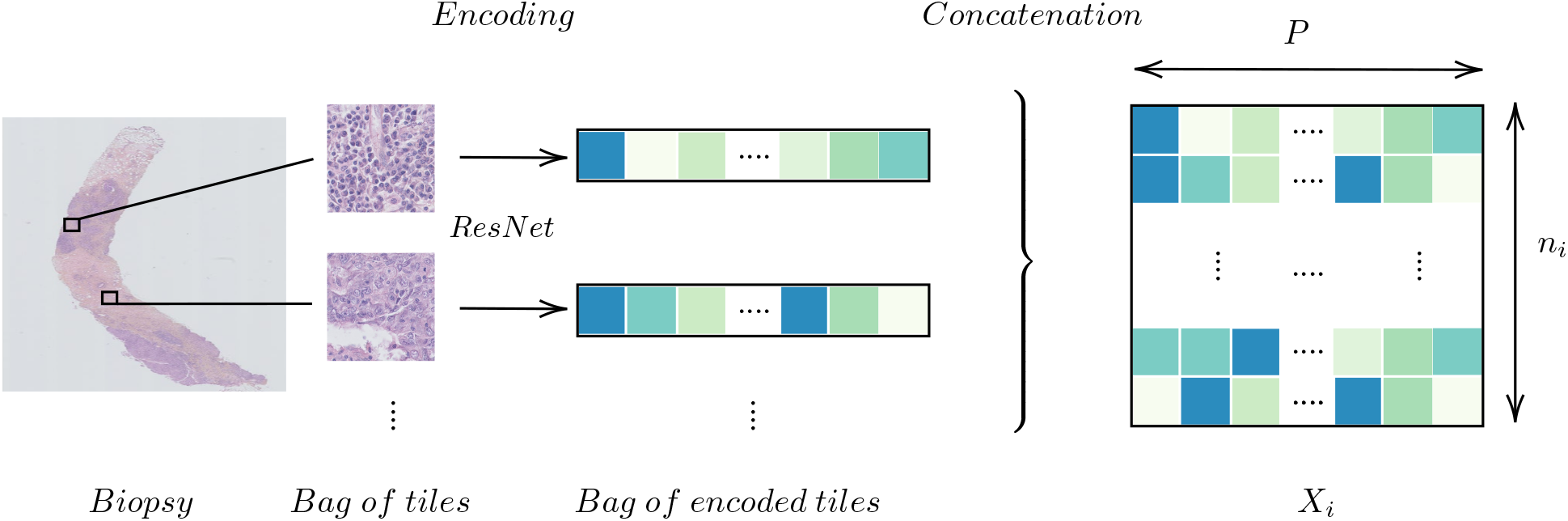
Encoding a biopsy

In Table 1 we show the average number of tiles, 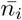 and variance at different magnification factors: highest resolution i.e. no down-sampling (2^0^ = 1), down-sampling by a factor 2^1^ and by a factor 2^2^ = 4.

**Table 1:** Mean number of tiles

| Down-sampling factor | $\bar{n}_i$: Mean number of tiles | $\sum_i n_i$ |
| --- | --- | --- |
| $2^0$ | $11\,186 \pm 6\,983$ | 3 758 389 |
| $2^1$ | $2\,757 \pm 1\,757$ | 926 409 |
| $2^2$ | $628 \pm 403$ | 211 077 |

The size of the data remains relatively large even after this reduction. We further reduce the size of the tile encoding with a PCA [50], and project each tile encoding into a space approximately 10× smaller. By keeping 256 components, we keep 93.2% at 2^0^, 94.0% at 2^1^ and 94.3% at a magnification factor of 2^2^ of the explained variance.

#### 2.3.3 Mathematical framework

The data set will be denoted by 𝒟 = (*X*_*i*_, *Y*_*i*_)_*i*∈ ⟦1,*N* ⟧_, and every item indexed by *i* in 𝒟 is a joint variable (*X*_*i*_, *Y*_*i*_) where *N* is the size of the data set, *X*_*i*_ is the input sample and *Y*_*i*_ the corresponding label. As described in the previous section, each tissue is represented by a bag of tiles of variable sizes, in particular 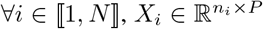 and *Y*_*i*_ ∈ {0, 1} for task (1) or (2). This is simply a multiple instance learning framework, and such a framework has already been implemented for histopathological data [28,51–53]. We simplify this framework by setting ∀*i* ∈ *i* ∈ ⟦1; *N* ⟧, *n*_*i*_ = *n*_*MF*_ ^2^ which is set accordingly to the chosen magnification factor. For a given sample *i*, if *n*_*i*_ *> n*_*MF*_ we down-sample *X*_*i*_, otherwise we up-sample *X*_*i*_ to the correct size. We evaluate our models by using the Area Under the Curve of the Receiver Operating Characteristic for measuring the performances in our two binary settings.

### 2.4 Neural Network architectures

Today, DNN models for WSI classification usually consist in 3 steps: starting from encodings that are usually provided by pre-trained networks, a reduction layer might be applied, followed by an aggregation step that computes a slide level representation from the tile level representations and a final module that maps the slide level representation to the output variable.

In Figure 3, we show a basic example for such an architecture along these lines, with the three algorithmic blocks highlighted in gray. At the tile-level computation, we use 1D convolutions to transform the input encodings *X*_*i*_ into a more compact representation 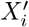. The tile representations 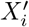 are then summarized by a pooling layer, providing us with the biopsy section profile *Z*_*i*_. For this, we can use standard pooling layer such as average pooling to quantify the abundance of specific tile patterns, or more complex, attention-based pooling, such as WELDON [54]. Finally, from *Z*_*i*_, the slide variable is predicted.

**Figure 3:**
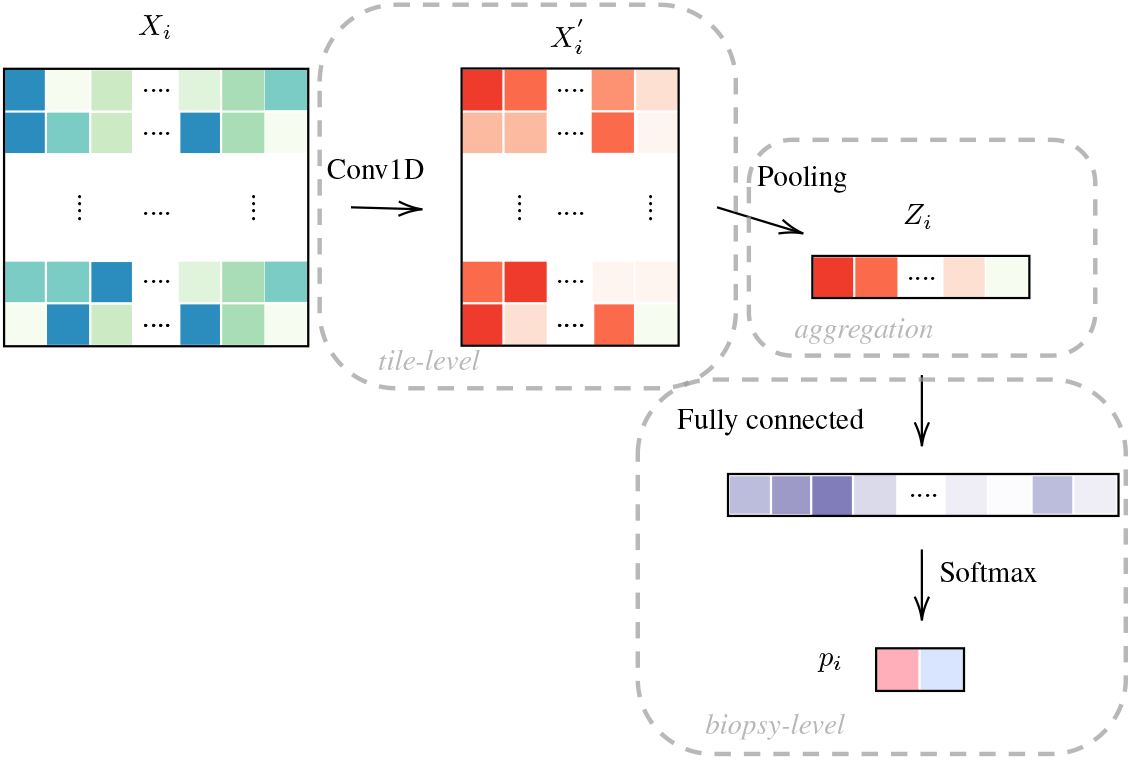
Model OneOne: tile encodings *X*_*i*_ from a pretrained network are projected to a more compact representations 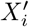 and then aggregated to build the slide representation *Z*_*i*_, which is then used for prediction. Same colors indicate identical dimensions.

In this article, we test several encoding and agglomeration strategies which are explained in the next sections.

#### 2.4.1 Encoding projection

In Figure 3, the baseline approach is illustrated (OneOne), with a 1D convolution for the tile level encoding and one fully connected layer at the slide level. In Figure 4, we test a deeper architecture for the encoding projections, consisting in 3 consecutive 1D convolutions, including bottleneck layers (depicted in orange), according to best practice in deep learning [55].

**Figure 4:**
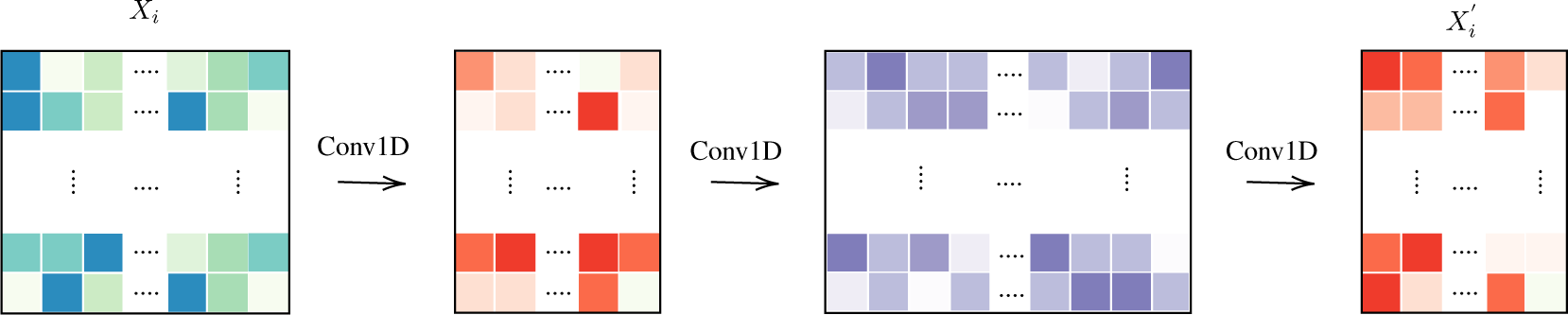
The computation: Three layers

Furthermore, we also experiment with skip connections by concatenating the first tile representations to the final representation 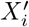 and by concatenating *Z*_*i*_ prior to the final softmax. We name this structure ThreeTwoSkip and illustrate it in Figure 5.

**Figure 5:**
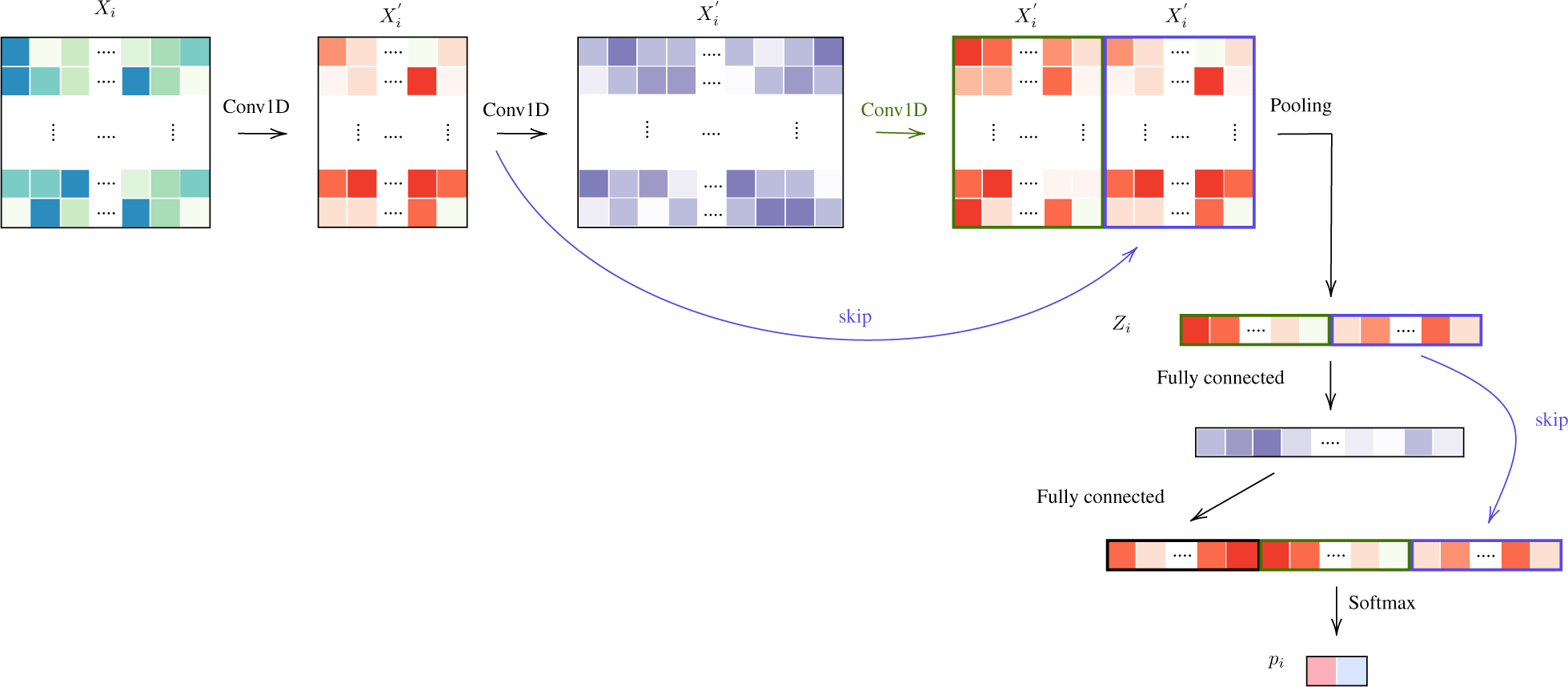
ThreeTwoSkip

#### 2.4.2 Pooling layers

In terms of pooling layers, we experiment with: average pooling shown in Figure 6.A, WELDON [54] shown in Figure 6.B, a modified version of WELDON shown in Figure 6.C and the concatenation of the first and the third is named WELDON-C (for context). The DNN that uses WELDON-C will be named CONAN^3^.

**Figure 6:**
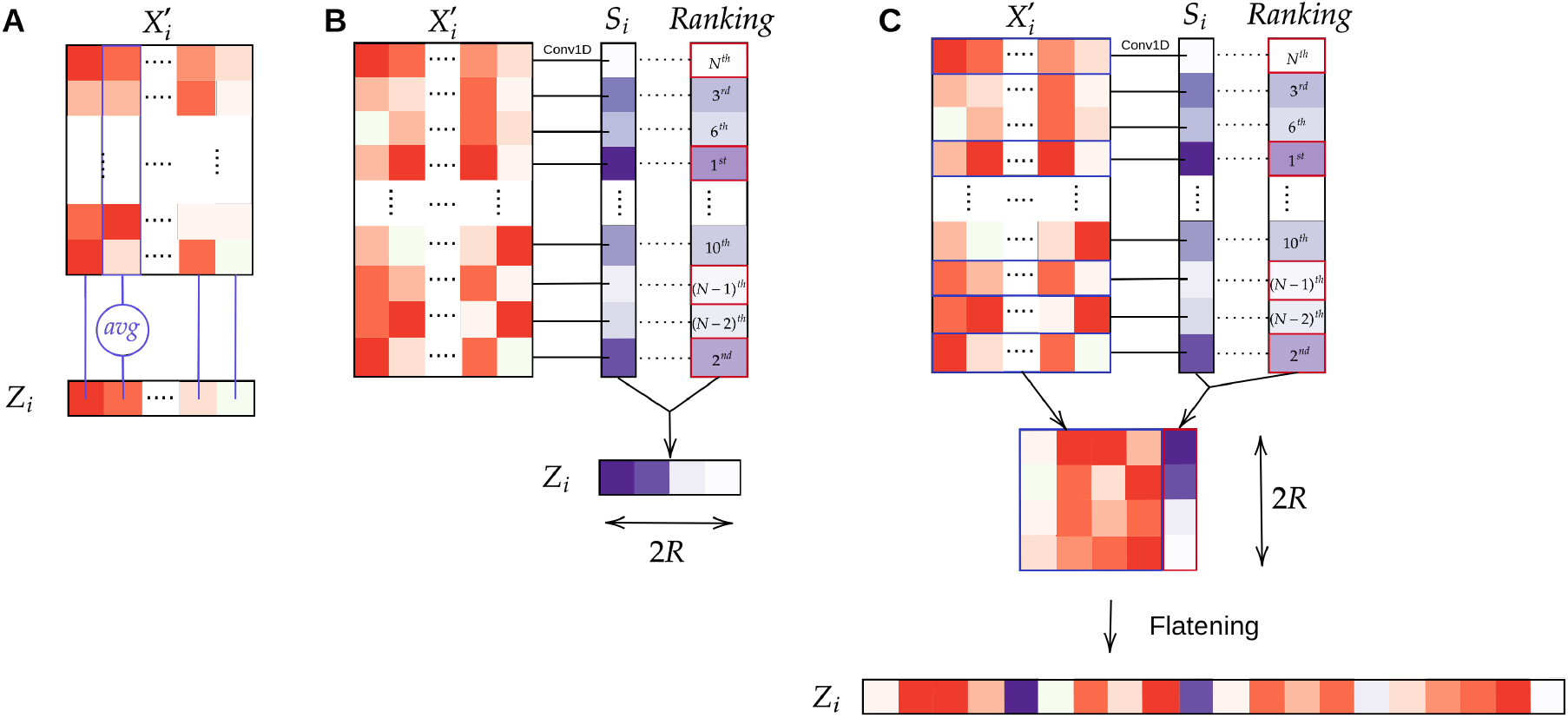
Aggregation layers: (**A**) Average pooling; (**B**) WELDON pooling; (**C**) WELDON-C pooling which is WELDON concatenated with previous tile encoding.

The WELDON pooling is an attention-based layer which filters tiles based on a 1D convolution score. In particular, it retains the top and lowest *R* ∈ ℕ^*^ achieving scores as *Z*_*i*_. This architecture has shown excellent results for specific problems where the biological evidence lies in the detection of one type of specific tiles, like cancer regions [28]. The method however suffers from identifiability issue, i.e. the model can not differentiate between two tiles achieving high or low score. In addition, the agglomeration strategy seems less promising in cases where the information resides in the percentage of tiles of a certain type. By providing a context in which a tile was selected, we allow the model to better differentiate between the selected tiles, thus allowing different tiles with different meanings to be selected, this can be particularly efficient when relevant information is based on different tile patterns.

We recap all the tested models in Table 2.

**Table 2:**
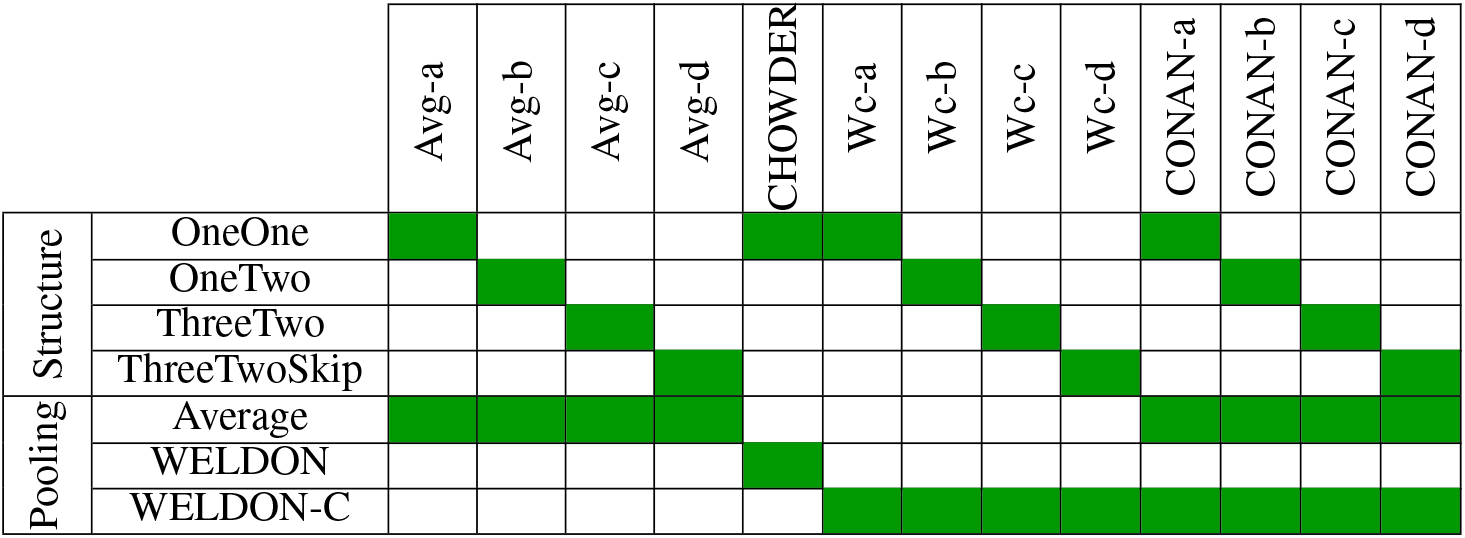
Model architecture and pooling strategy, as described in the text. CHOWDER has been published previously [28].

#### 2.4.3 Model tuning

We perform a random grid search for most parameters and only in suitable ranges. For the learning rate and weight decay we perform a random log sampling for a random scale associated to a random digit. We range from a scale of 10^−6^ to 10^−3^ for the learning rate and from 10^−4^ to 10^−1^ for the weight decay. We randomly sample a drop out from a uniform 𝒰 _[0:0.4]_. We randomly sample a bottleneck layer size from the following list [8, 32, 64] and the size of the larger representations are randomly sampled from [64, 128]

## 3 Results

### 3.1 Simulation results

#### 3.1.1 High Performance variability in DNN training

In Figure 7.A, we show the average variance of our model with increasing standard deviation *σ*. In particular, for each *σ*, we generate 100 simulated dataset with a standard deviation of *σ* and train 1000 DNN on the same data and with the same HP. As this is simulated data, we evaluate the performance of each training on a large independently simulated test set instead of using the outer CV loop [32]. We found that setting the learning rate to 1.10^−4^, the weight decay to 5.10^−3^ and drop out to 0.4 tend to always return reasonable results for our simulation setting.

**Figure 7:**
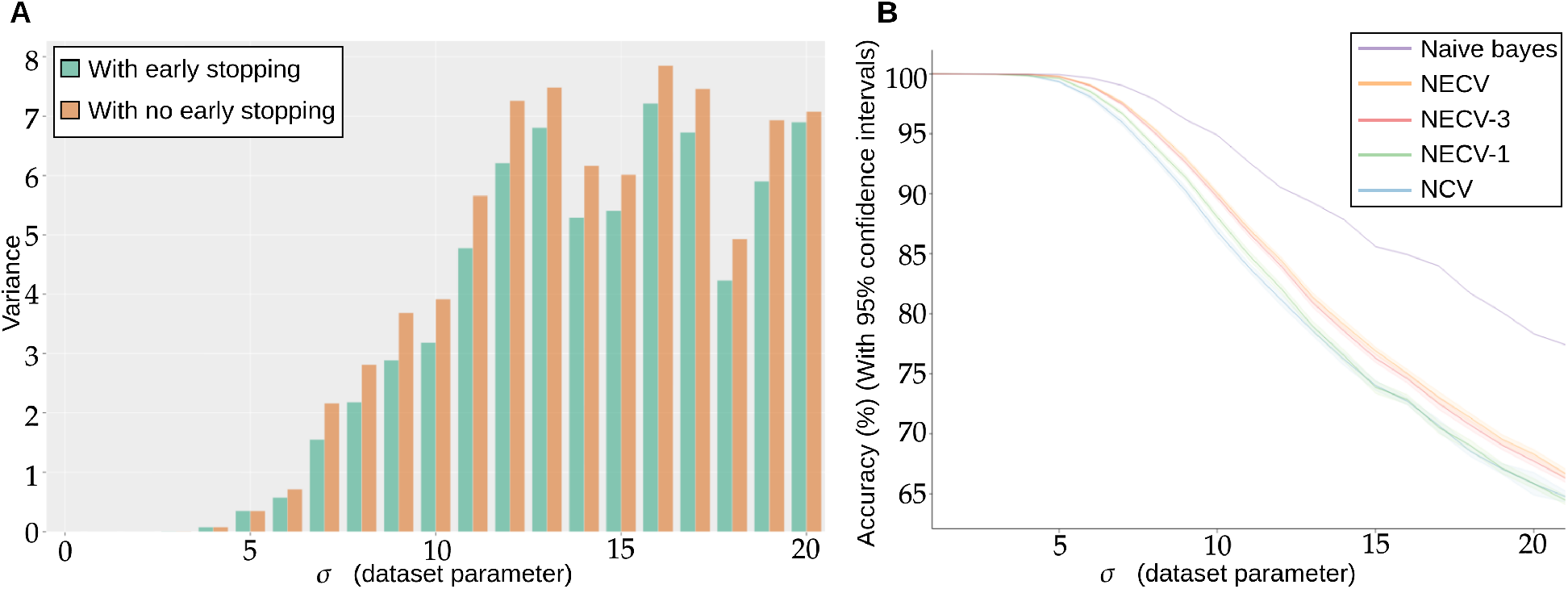
(**A**) Repeated training, with and without early stopping: (**A**) Performance Variance, as measured on an independently simulated test set, (**B**) Comparison of models trained with NCV and NECV. Naive Bayes is a theoretical upper limit of the achievable performance, by construction of the simulation. Accuracies are shown with 95% confidence intervals.

As the standard deviation *σ* of the simulated data increases, we expect more overlapping between our two classes and naturally, the classification accuracy decreases. For lower *σ*, regardless of using early stopping or not, the models reaches perfect scores.

In Figure 7, we observe that the more difficult the problem (larger *σ*), the lower the accuracy, but also the larger the variance: not only do we predict less well, but also does the performance variation increase, such that by retraining a model with the same hyperparameters is not guaranteed at all to provide a model with similar performance. We also see from Figure 7.A that early stopping alleviates this problem and consistently reduces the variance in performance, in particular for higher *σ*.

#### 3.1.2 NCV leads to under-performing models

We compare the performance of the proposed validation procedure to NCV with the number splits *k*_*f*_ = 5 in Figure 7.B with 95% confidence intervals around the estimator. On the *x*-axis we have the standard deviation *σ* of the simulated data and on the *y*-axis the averaged corresponding performance of NCV or NECV. For each *σ*, we collect 40 estimators with Algorithm 1 and 3. Next, we compare several NECV strategies: NECV-1 (green curve), where we keep the best model, NECV-3 (red curve), where we keep the top 3 models and NECV (orange curve), where we keep all 5 models.

We first notice that the NCV curve is lower or equal to any of the NECV curves. The best performing model is NECV – i.e. the average of all selected models from the inner CV. In particular NECV has a higher Accuracy than NCV by at least 2%. We conclude that retraining the model without an outer validation score leads to lower overall performance and early stopping is a very useful regularization technique for small sample size problems.

### 3.2 Prediction of response to neoadjuvant chemotherapy

We next applied the different architectures detailed in section 2.4 and summarized in table 2 to the problem of the prediction of response to neoadjuvant chemotherapy in TNBC. We tested 3 different image resolutions (0, 1, 2), 0 being the highest resolution. In order to get realistic estimations of the performance, while using early stopping, we perform the validation proposed in Section 2.1. In Figure 8.A and 8.B we show the average AUC ROC performance on the residual and prognostic prediction tasks, for all methods shown in Table 2 and for all resolution levels.

**Figure 8:**
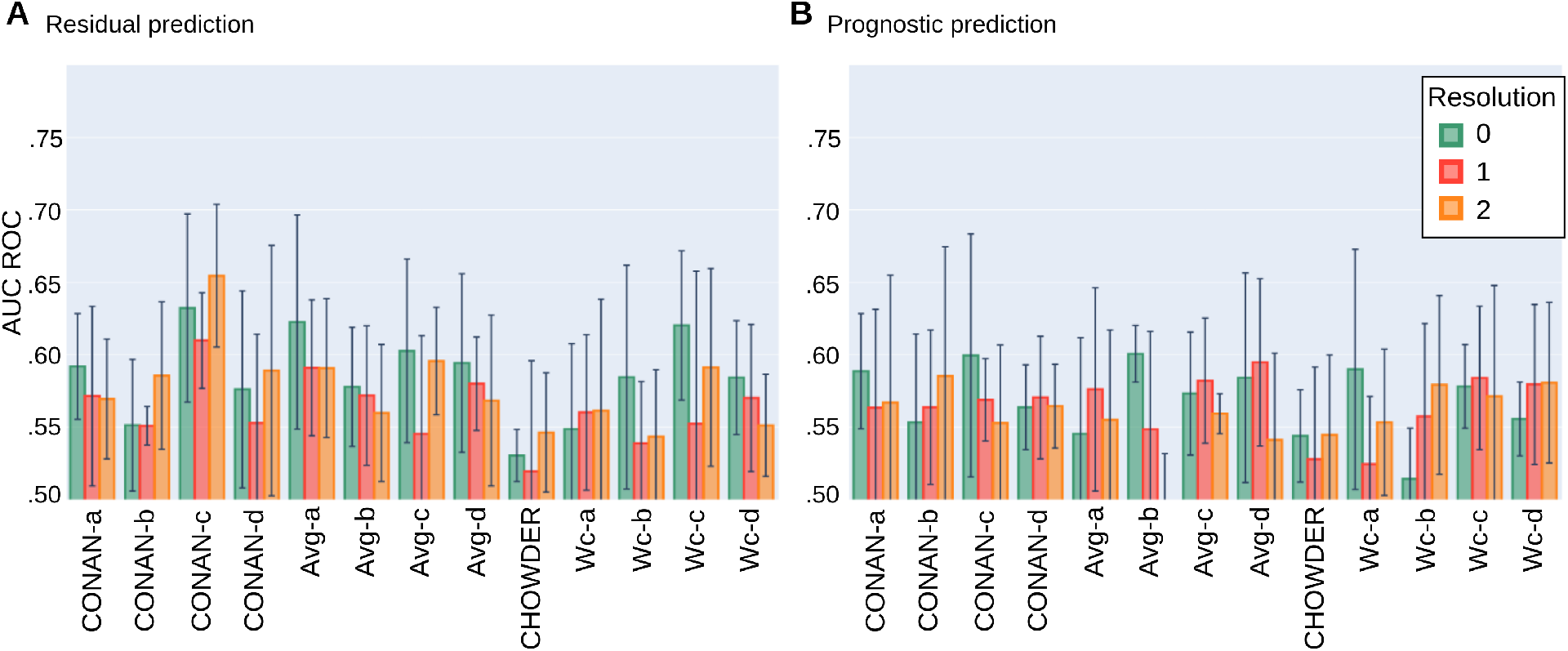
Average AUC ROC performance with standard deviation on 5 fold NECV on the task of predicting: (**A**) residual cancer and (**B**) patient prognostic in TNBC.

For the task of predicting the residual cancer, the best performing model would be the *CONAN-c* model at resolution 2 with an AUC of 0.654 ± 0.049. Others model’s performance range in between 0.55 and 0.60 of AUC with higher standard deviations. Models at resolution 0 seem to generally achieve higher scores then those at lower resolutions. Model architecture *c* seem to be better suited for this task than the others. The Average concatenated to WELDON-C pooling seems to perform slightly better then the rest. The method CHOWDER which gave excellent results on CAMELYON for cancer detection [28] and which has been a state-of-the-art solution in the field under-performs on our dataset for response prediction.

For the task of predicting the patient prognostic, the best performing model would be the *Avg-b* model at resolution 0 with an AUC of 0.601 ± 0.019. *CONAN-c* at resolution 2 performs similarly but with a much higher standard deviation. Neither resolution, nor model architecture and pooling layer seem to unanimously be better then the others. However, CHOWDER under-performs compared to the other proposed methods.

## 4 Discussion

In this study, we set out to predict the response to neoadjuvant chemotherapy in TNBC from biopsies taken before treatment. A system that would allow to predict this response with high accuracy could help identifying patients with no or little benefit of the treatment and therefore spare them the heavy burden of the therapy.

From a methodological point of view, this is particularly challenging for three reasons: first, we do not know to which extent the relevant information is actually present in the image data. In addition, even if the relevant information is contained in the slide, the complexity of the related patterns is unclear. Second, biopsies only capture a part of the relevant information, as they are only a localized sample of the tumor. Third, as this is a project regarding a specific subtype, the cohort is relatively small, unlike many pan-cancer cohorts used in large Computational Pathology projects [17].

In order to solve this problem, we have developed the model *CONAN*, that combines the power of selecting *K* tiles (top and bottom), but keeps both the ranking scores and the full tile descriptions to build the slide representation. We have compared this model with a number of different architectures, and achieved an AUC of 0.65.

We also tackled an important problem of model selection with cross validation, a crucial step in particular for small datasets. We found that the retraining step in classical Nested Cross Validation can lead to lower performances for small *N*, because the training is highly variable, and a network retrained with the optimal set of hyperparameters is not guaranteed at all to be optimal itself. We therefore have proposed a new cross validation procedure relying on ensembling rather than retraining, and thus allowing to use early stopping as a regularization method.

Nevertheless, we must conclude that the prediction of treatment response is probably one of the hardest problems in Computational Pathology, and that even though we see that there is some degree of predictability, the results still seem far from clinical applicability. Clearly, we need more data to tackle this challenging question. But it is also likely that even with much more data, AUCs will not reach very high levels by looking at biopsies alone. A promising avenue would therefore be to use other kinds of data in addition to histopathology data.

## Code availability

In addition to the methodological developments and in the spirit of reproducible research, we make the code for all experiments publicly available in the following github repository https://github.com/PeterJackNaylor/AutomaticWSI. The code was mostly written using Python3, Keras [56] and Nextflow [57].

## Acknowledgment

The authors would like to thank André Nicolas from the PathEx platform, Service the Pathologie, Institut Curie, Paris, where the images have been acquired. We would also like to thank Ligue National Contre le Cancer for funding the P.h.D of Dr. Naylor. Furthermore, this work was supported by the French government under management of Agence Nationale de la Recherche as part of the “Investissements d’avenir” program, reference ANR-19-P3IA-0001 (PRAIRIE 3IA Institute).

## Footnotes

2 For Magnification Factor

3 Context cOncatenated tile selection NeurAl Network

